# Return of a lost structure in the evolution of felid dentition revisited: A DevoEvo perspective on the irreversibility of evolution

**DOI:** 10.1101/2021.02.04.429820

**Authors:** Vincent J. Lynch

## Abstract

There is a longstanding interest in whether the loss of complex characters is reversible (so-called “Dollo’s law”). Reevolution has been suggested for numerous traits but among the first was Kurtén (1963), who proposed that the presence of the second lower molar (M_2_) of the Eurasian lynx (*Lynx lynx*) was a violation of Dollo’s law because all other Felids lack M_2_. While an early and often cited example for the reevolution of a complex trait, Kurtén (1963) and Werdelin (1987) used an *ad hoc* parsimony argument to support their proposition that M_2_ reevolved in Eurasian lynx. Here I revisit the evidence that M_2_ reevolved in Eurasian lynx using explicit parsimony and maximum likelihood models of character evolution and find strong evidence that Kurtén (1963) and Werdelin (1987) were correct – M_2_ reevolved in Eurasian lynx. Next, I explore the developmental mechanisms which may explain this violation of Dollo’s law and suggest that the reevolution of lost complex traits may arise from the reevolution of cis-regulatory elements and protein-protein interactions, which have a longer half-life after silencing that protein coding genes. Finally, I present a model developmental model to explain the reevolution M_2_ in Eurasian lynx.

## Introduction

There is a growing interest and renewed debate in whether the loss of complex characters is reversible. Reevolution of complex characters has been suggested for eyes in ostracods (Dingle, 2003; Hunt, 2007; Oakley, 2003), spiders (West-Ebehard, 2003), and snakes (Coates and Rutta, 2000; Laurent, 1983), ocelli in cave crickets (Desutter-Grandcolas, 1993), wings in water spiders (Anderson, 1997), fig wasps (Whiting and Whiting, 2004) and stick insects (Whiting et al., 2003), digits in lizards (Brandley et al., 2008; Kohlsdorf and Wagner, 2006), developmental stages in salamanders (Chippindale et al., 2004; Mueller et al., 2004) and frogs (Wiens et al., 2007), teeth in lynx (Kurtén, 1963) and marsupial frogs (Wiens, 2011), thigh muscles in birds (Raikow, 1975), and oviparity in a clade of otherwise viviparous snakes (Lynch and Wagner, 2010), among many others.

Kurtén (1963) described one of the first examples of the reevolution of a lost complex character—the second lower molar (M_2_) of the Eurasian lynx (*Lynx lynx*). Unlike all other extant felids, which lack M_2_, this molar is present in 3.4-27% of Eurasian lynx populations (**Figure 1**). Using an *ad hoc* parsimony argument, Kurtén suggested that M_2_ recently reevolved in the Eurasian lynx in association with the reevolution of the metaconid-talonid complex of the first lower molar (M_1_). This apparent break from Dollo’s law was analyzed in a more phylogenetic framework by Werdelin (1987), who showed that M_2_ was absent from other species of extant and extinct Lynx; following Kurtén, Werdelin proposed that the presence of M_2_ was a true reversal. While the presence of M_2_ in *Lynx lynx* and its implications for Dollo’s law were the subject of some debate (Kvam 1985), this example has become one of the most citied cases against strict enforcement of Dollo’s law. However, Kurtén’s proposal that M_2_ reevolved in the Eurasian lynx has never been explicitly tested.

**Figure 1.**
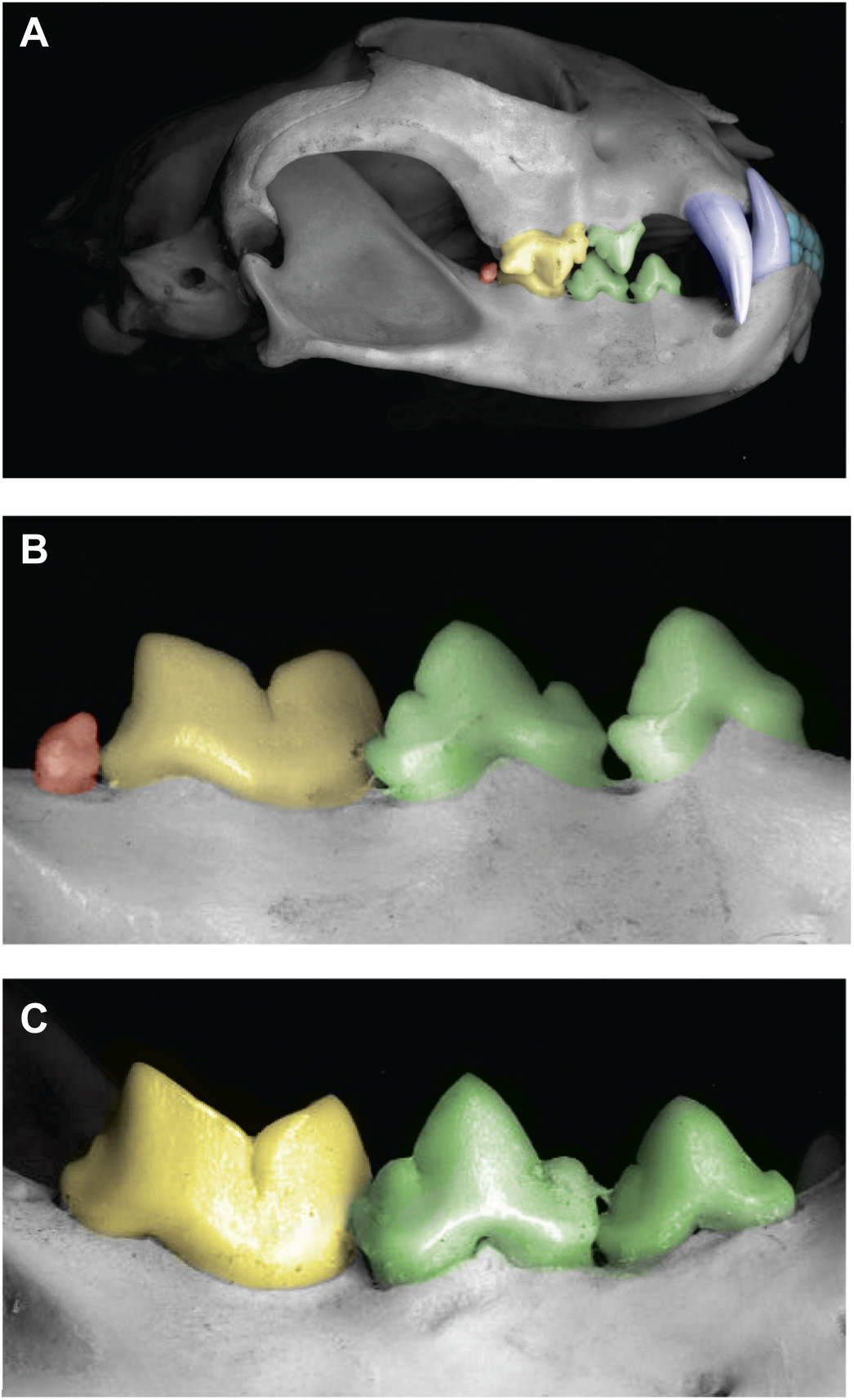
Dental variation in the Eurasian lynx (*Lynx lynx*). **(A)** Articulated skull, with incisors colored light blue, canines dark blue, premolars green, the first molar (M_1_) yellow, and the second molar (M_2_) orange. **(B)** Close up of a mandible with M_2_ colored orange. **(C)** Close up of a mandible without M_2_ colored.

In this paper I reanalyze the evidence for the reevolution of the second lower molar (M_2_) of the Eurasian lynx using explicit models of character evolution to evaluate Kurtén’s proposal that Dollo’s law was broken. First, I perform a molecular phylogenetic analysis of extant felids based on a published dataset of thirty nuclear and eight mitochondrial genes for 38 species of felids, including genera from each of the felid major lineages, plus seven feliform groups closely related to the Felidae to generate a phylogeny on which to test for revolution of M_2_ in Lynx. Using this well-supported phylogeny, I reconstruct the phylogenetic history of the second molar in extant cats using parsimony- and likelihood-based models of character evolution and present strong statistical evidence that M_2_ reevolved in the Eurasian lynx, reaffirming Kurtén’s hypothesis. Finally I present a statistical model to explain the potential developmental genetic basis of reevolution, and simple developmental model to explain the mechanisms that may underlie the reappearance of M_2_ in the Eurasian lynx.

## Materials and Methods

### Phylogenetic analysis

We used the Johnson et al. (2006) dataset to generate a phylogeny for character reconstructions, which included 45 species and 23,920 nucleotide sites from thirty nuclear loci, *APP, CALB, CHRNA, CLU, CMA, DGKG2, FES, GATA, GHR, GNAZ, GNB, HK1, NCL, PNOC, RAG2, RASA, SILV, TCP, TTR, ALAS, ATP7A, IL2RG, PLP, ZFX, SMCY, SRY3, SRY5, SRY, UBEY, ZFY*, and eight mitochondrial loci *16s, ATP8, CYTB, ND1, ND2, ND4, ND5*, and *t-RNAs*. Alignments were the same as in the original study, and regions of ambiguous alignment were excluded from further analysis. Bayesian phylogenetic analyses were performed with MR.BAYES v3.1 (Huelsenbeck and Ronquist, 2001) using a partitioning scheme that applied a separate GTR+Γ substitution models to each gene and unlinking model parameters across each partition; more complex partitioning schemes, such as those dividing genes into codon positions, were evaluated but generally were too computationally complex and failed to converge. Two independent Bayesian analyses were run for 10,000,000 generations each with four chains sampled every 1000 generations and a burnin of 2,500 trees. Run progress was visually checked with TRACER v1.4 by plotting the log-likelihoods of sampled generations, and the stability of parameter estimates (chain convergence) checked by ensuring that the standard deviation of split likelihood frequencies was below 0.01 while the potential scale reduction factor (PSRF) was close to 1.0 for all parameters.

### Character Evolution

Parsimony and likelihood reconstructions of character evolution were performed with MESQUITE v2.01 (Maddison and Maddison, 2009). The rates of forward (*q*_*01*_) and reverse transitions (*q*_*10*_) under an asymmetrical likelihood model with separate rates for forward and reverse transitions estimated directly from the data (rMk2) and compared to an irreversible model (iMk2) that constrained the reverse rate to zero (*q*_*10*_=0). Because recent studies indicate that character associated changes in diversification rate can lead to erroneous rejection of irreversible models (Goldberg and Igić, 2008), we tested for character associated diversification using the BiSSE module (Maddison et al., 2007) implemented in MESQUITE v2.01. For both Markov and BiSSE models a stationary or “uninformative” prior was used to infer the state of the root node under reversible models while irreversible models fixed the state of the root node at 0 in the BiSSE analysis. The tree for character reconstructions was the consensus derived from the Bayesian analysis transformed into ultrametric tree with an arbitrary root age of 100 using the penalized likelihood method implemented in R8S v1.7 (Sanderson, 2003).

Parsimony based trait reconstructions were analyzed as unordered (reversible) and imposing the ‘Dollo’ model in MacClade. To compare the statistical support for different parsimony models we used the ‘no common mechanism’ model (NCM) of Tuffley and Steel (1997) to transform tree length into likelihoods. Calculation of the NCM model score followed equation 42 of Tuffley and Steel (1997),

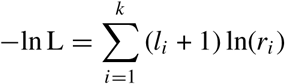

where *l*_*i*_ is the parsimony cost of the reconstruction given the tree, and *r*_*i*_ is the number of states in character *k*; because only a single character is considered here, the NCM model is equivalent to a scaling of the parsimony tree length into -log likelihoods.

### Probabilistic models of escape from deleterious mutations

Following Marshall, Raff, and Raff (1994), I calculated the probability that genes retain function during periods of “silencing”. These authors proposed that the probability a gene (or regulatory element) retains a functional sequence in the absence of selection is

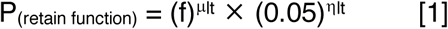

where *f* is the probability that a nucleotide substitution is not inactivating, *μ* is the per nucleotide substitution rate in millions of years (Myr), *η* is the rate of frameshifts (Myr), *l* is the length of the sequence, *t* is time (Myr) and 0.05 indicates that frameshifts in all but the last (most C-terminal) 5% of a gene are deleterious. Using empirical data, Marshall, Raff and Raff (1994) calculated a liberal *f* to be 0.7, and a moderate *μ* and *η* to be 0.005 and 0.093, respectively. Here we extend the Marshall, Raff, and Raff (1994) model from large protein coding genes to include sequences of size that are typical for both cis-regulatory elements and individual protein-protein interaction sites in proteins.

### Results and Discussion

Unlike all other extant felids, 3.4-27% of Eurasian lynx populations have a second lower molar (M_2_; **Figure 1**). Using an *ad hoc* parsimony argument, Kurtén (1953,1963) suggested that M_2_ recently reevolved in the Eurasian lynx, a proposition supported by Werdelin (1987) using a more phylogenetic framework. Here we reevaluated the evidence supporting the reevolution of M_2_ in the Eurasian lynx using a large, well-supported, phylogeny and contemporary models of character evolution.

### Phylogeny of the Felidae

The Bayesian phylogeny inferred for the felids is very similar to those previously estimated using molecular data, and is nearly identical to the tree inferred by Johnson et al. (2006). We found that the majority nodes in the tree were supported by Bayesian posterior probabilities (BPP) of 1.0, with the exception being three nodes in the domestic cat lineage with BPP of 0.56, 0.71, and 0.91 (**Figure 2**). Although the molecular dataset used to infer the phylogeny (**Figure**

**Figure 2.**
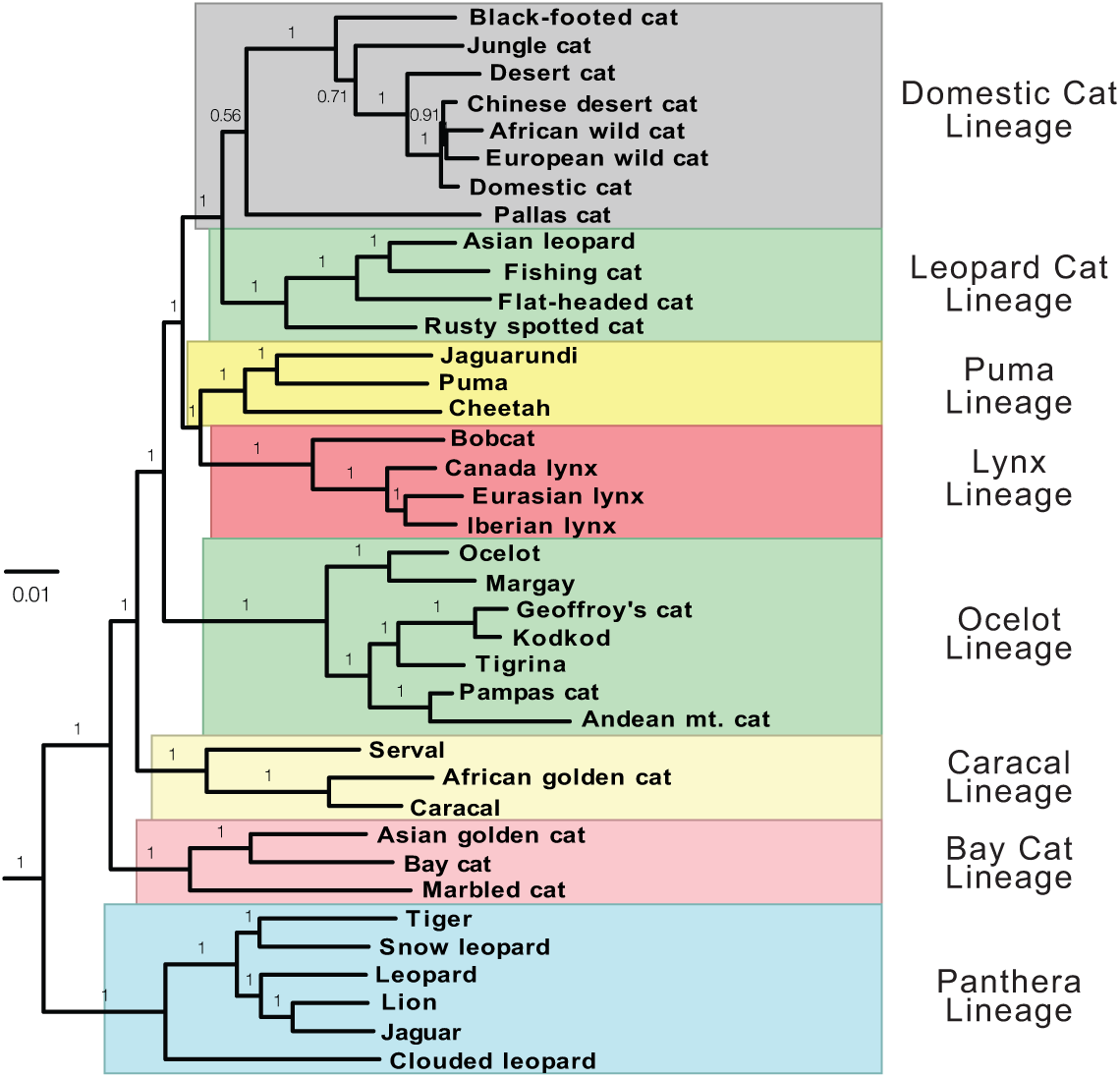
Felid phylogeny. Bayesian phylogeny of extant cats based on a concatenated alignment including 45 species and 23,920 nucleotide sites from 30 nuclear loci and eight mitochondrial loci. Branch lengths are proportional to substitutions per site, Bayesian Posterior Probabilities (BPP) for each branch. Major lineages are named and highlighted.

**2)** is identical to the one by used Johnson et al. (2006), there are two differences between our tree topologies. We find that the Pallas cat is placed as the most basal lineage in the domestic cat clade, albeit with very poor support (BPP: 0.56), while Johnson et al. (2006) found the Pallas cat to be sister to the leopard lineage also with relatively poor support (BPP: 0.83). Second, we find that the puma lineage is placed as the sister clade to the lynxes (BPP: 1.0), rather than as sister to the leopard+domestic cat lineage as did Johnson et al. (2006), again with very poor support (BPP<0.50).

A preliminary analysis indicates that these different topologies result from different data partitioning schemes; Johnson et al. (2006) used as single GTR+Γ+I model in their combined nuclear/mitochondrial Bayesian analysis while I used separate GTR+Γ+I models for each codon position of protein coding genes and each stem and loop of RNA genes. Penalized likelihood analysis using our phylogeny and the 15 fossil calibrations used by Johnson et al. (2006) gives essentially the same divergence times as the original study and interested readers are referred to Johnson et al. (2006) for the systematic and biogeographic implications of felid phylogeny. We note, however, that these phylogenetic differences do not alter our inference of M_2_ loss and gain in felids.

### Reevolution of M_2_ in Eurasian lynx: Parsimony

To infer whether explicit models of character evolution support the inference of M_2_ reevolution, I reconstructed the evolutionary history of M_2_ in cats using the well-supported molecular phylogeny plus the extinct Issoire lynx (*L. issiodorensis*) of felids and parsimony reconstruction coding the Eurasian lynx as polymorphic for the presence of M_2_. I found that a reversible model required only two steps (one basal loss in the stem-lineage of the Felidae, one regain in the Eurasian Lynx), while the irreversible ‘Dollo’ model, which disallows regains and thus imposes multiple losses of M_2_, required 10 steps (**Table 1** and **Figure 3**). Comparison of NCM likelihoods between these two models indicates that there is considerably less support for the irreversible ‘Dollo’ model (-LnL = −7.62) than for the reversible model (-LnL = −2.08), indicating that there is essentially no support for the ‘Dollo’ model (ΔAIC = 11.09; **Table 1**).

**Table 1.**
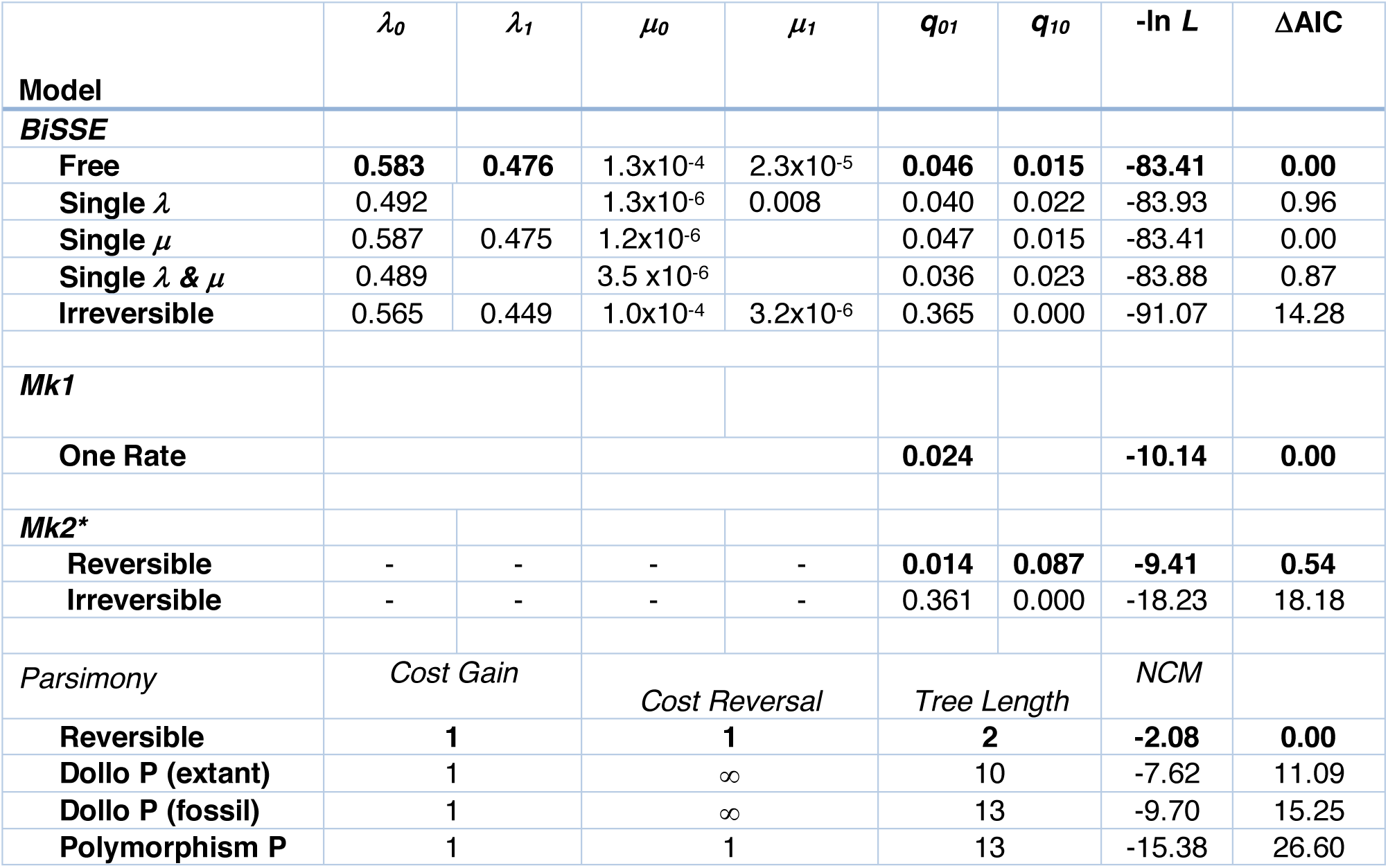
Model comparisons testing the irreversibility of molar loss. Speciation (***λ***), extinction (***μ***) and character transition (***q***) rate estimates are reported based on the time calibrated tree with an arbritary root age of 100. The likelihood of each model (ln *L*) is shown, followed by the ΔAICs; the best models are in bold. Note that the BiSSE model does not support trait dependent diversification rates, therefore our conclusions on reversibility are based on the Mk2 model. BiSSE and parsimony results are shown for comparison only. The No Common Mechanism (NCM) score is a scaling of the parsimony tree length into -log likelihoods and is shown for comparison to the models. Note that the parsimony model includes the extinct Issoire lynx (*L. issiodorensis*) that is not included in either the BiSSE or Mk models.

**Figure 3.**
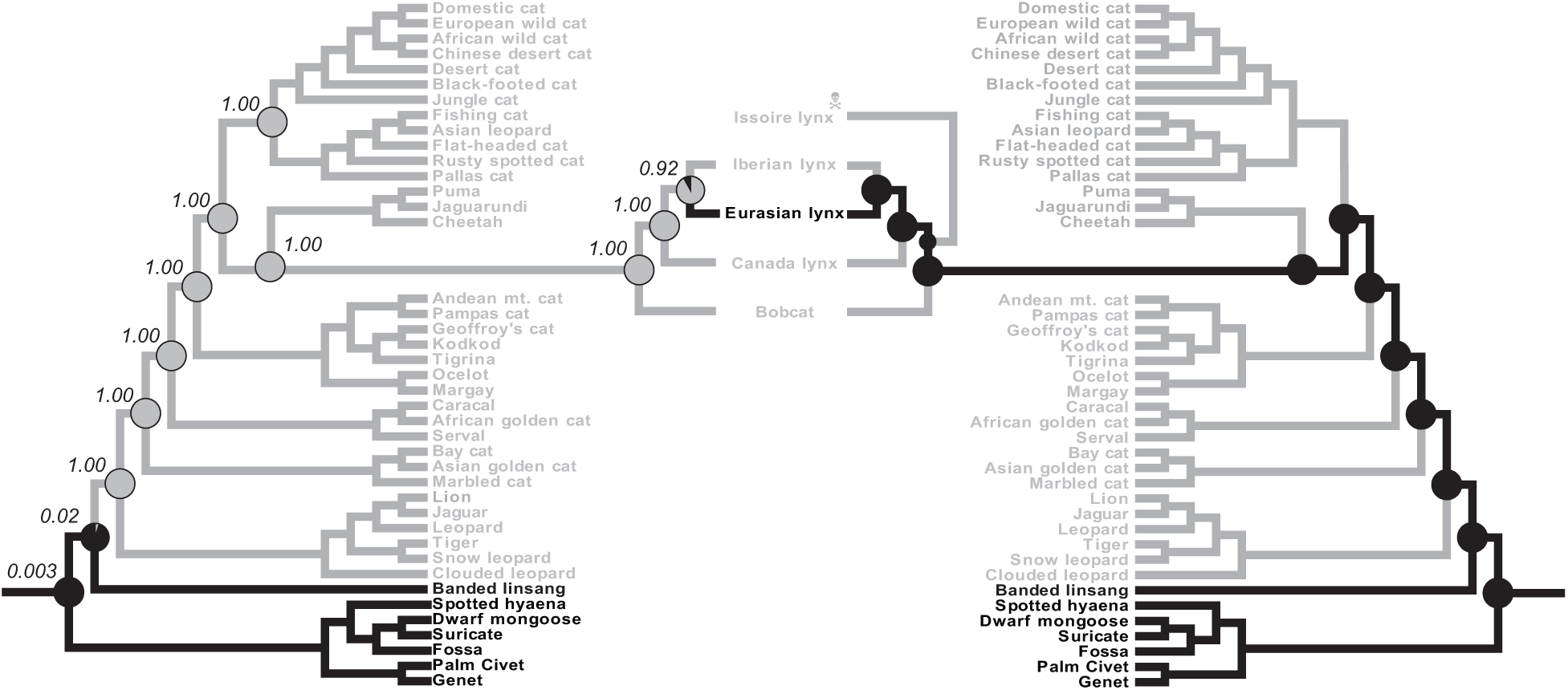
Phylogenetic support for the reevolution of M_2_ in Eurasian lynx. Left, Reconstruction of ancestral states under the reversible Mk2 model. The probability of M_2_ presence (black) and absence (grey) is shown at each ancestral node as a pie chart colored by the probability of each state. The probability of M_2_ absence is shown at each node. Branches are colored according to the reversible parsimony model. Right, Reconstruction of ancestral states under the irreversible Mk2 model. The probability of M_2_ presence (black) and absence (grey) is shown at each ancestral node as a pie chart colored by the probability of each state. Branches are colored according to ancestral states under an irreversible (Dollo) parsimony model.

### Reevolution of M_2_ in Eurasian lynx: Likelihood

I also used two maximum likelihood models to test the irreversibility of M_2_ loss: methods that compare an irreversible model to models that allow reversals without accounting for the effect of the character on rates of speciation and extinction (Mk2 models) and likelihood models that accommodate for the effect of a character on speciation and extinction rates (BiSSE models). All likelihood analysis were performed on the time calibrated tree, including only extant taxa.

I found that the irreversible Mk2 model (iMk2: -LnL = −18.23) was a significantly worse fit to the data than the reversible Mk2 model (rMk2: -LnL = −9.41), indicating there was no support the irreversibility of M_2_ loss (**Table 1**), i.e. a Dollo model (ΔAIC = 18.18). In addition, the rMk2 model was only a marginally better fit to the data than a one-rate likelihood model (Mk1: -LnL = - 10.14), suggesting that there was little gain in power by including two rates of character evolution (ΔAIC = 0.54). Reconstruction of the most recent common ancestor of the felids under both the rMk2 and Mk1 models indicated there was a high proportional likelihood it lacked M_2_ (0.98), similarly reconstruction of the most recent common ancestor of the *Lynx* clade and *Lynx lynx*-*Lynx pardinus* lineages indicated there was a high likelihood these ancestors lacked M_2_ (1.0 and 0.92, respectively; **Figure 3**).

Like parsimony, Mk models are prone to falsely reject irreversibility when a characters state influences diversification rates (Goldberg and Igić, 2008). To account for the possibility that M_2_ presence/absence altered diversification rates in felids, confounding the inference of reversibility in parsimony and Mk models, we used the BiSSE model. The ‘free’ BiSSE model (-LnL = −83.41), which estimates different speciation and extinction rates for species with states 0 and 1, was only 0.22 likelihood units better than a model constrained speciation rates to be shared between state 0 and 1 (ΔAIC = 0.96). Similarly, the ‘free’ model was only a slightly better fit to the data than models that constrained extinction rates (ΔAIC = 0.00) or both speciation and extinction rates (ΔAIC = 0.87) to be shared between state 0 and 1. Thus, there is essentially no support for M_2_-dependent differences in diversification rates (**Table 1**). A BiSSE model that constrained M_2_ loss to be irreversible (-LnL = −91.07), however, was a significantly worse fit to the data than the ‘free’ model (ΔAIC = 14.28; **Table 1**).

### Is there a grace period on Dollo’s law?

Dollo’s argument for irreversibility was not merely an empirical generalization from the facts of the history of life (phylogeny), rather for Dollo it was a “question of probabilities”; complex characters do not reevolve because this would require that the organism retrace, in exactly the same order, an extremely large number of steps (Gould 1970). Thus, in the words of Muller, the loss of complex characters is irreversible because of “the sheer statistical improbability, amounting to an impossibility, of evolution ever arriving at the same complex genic end-result twice” (cited in Gould 1970). A developmental evolutionary basis for understanding Dollo’s rational is that complex characters arise from the execution of complex developmental genetic regulatory networks; the absence of stabilizing selection will lead to mutational drift in the components of the network and their loss. Therefore, its unlikely that the genetic information for the development and function of the character can be maintained for long periods of time in the absence of the character (Bull and Charnov, 1985).

Using empirical data and a statistical model to assess the probability that silenced genes escape these kinds of deleterious mutations, Marshall, Raff and Raff (Marshall et al., 1994) estimated that reactivation of silenced genes was likely if the time of inactivation was on the order of 0.5-6 million years. In contrast, resurrecting long dormant genes (> 10 million years) was extremely unlikely because of the accumulation of multiple inactivating mutations unless gene function was maintained in other contexts. The survival of protein coding genes, however, may not be sufficient to explain the reemergence of dormant developmental programs because the development of complex characters also depends on the correct spatial and temporal expression of those genes and their interactions with other genes. For example, many multifunctional proteins have tissue specific *cis*-regulatory elements that direct their appropriate expression (Davidson, 2006) and specific protein-protein interaction motifs that mediate their biochemical functions (Neduva and Russell, 2005, 2006).

I quantified the probability that these smaller *cis*-regulatory element and protein-protein interaction motif sized regions escape deleterious mutations following the model of Marshall, Raff and Raff (1994). I found that 100bp regions, which approximate the size of individual regulatory elements, such as enhancers, and small protein interaction domains, have a half life of 25 million years, while 10bp regions, corresponding to the size of individual transcription factor binding-sites and protein-protein interaction sites in proteins such as transcription factors, have a half life of around 250 million years (**Figure 4A**). In stark contrast, short (∼1kb), intermediate (∼3kb) and long (∼5kb) protein coding genes that been silenced only have 50% chance of surviving lethal mutations after 2.5, 0.85 and 0.5 million years (**Figure 4A**), respectively. These data indicate that reactivation of silenced enhancers, individual transcription factor binding-sites, and protein-protein interaction motifs can occur after a much longer time than an entire protein-coding gene.

**Figure 4.**
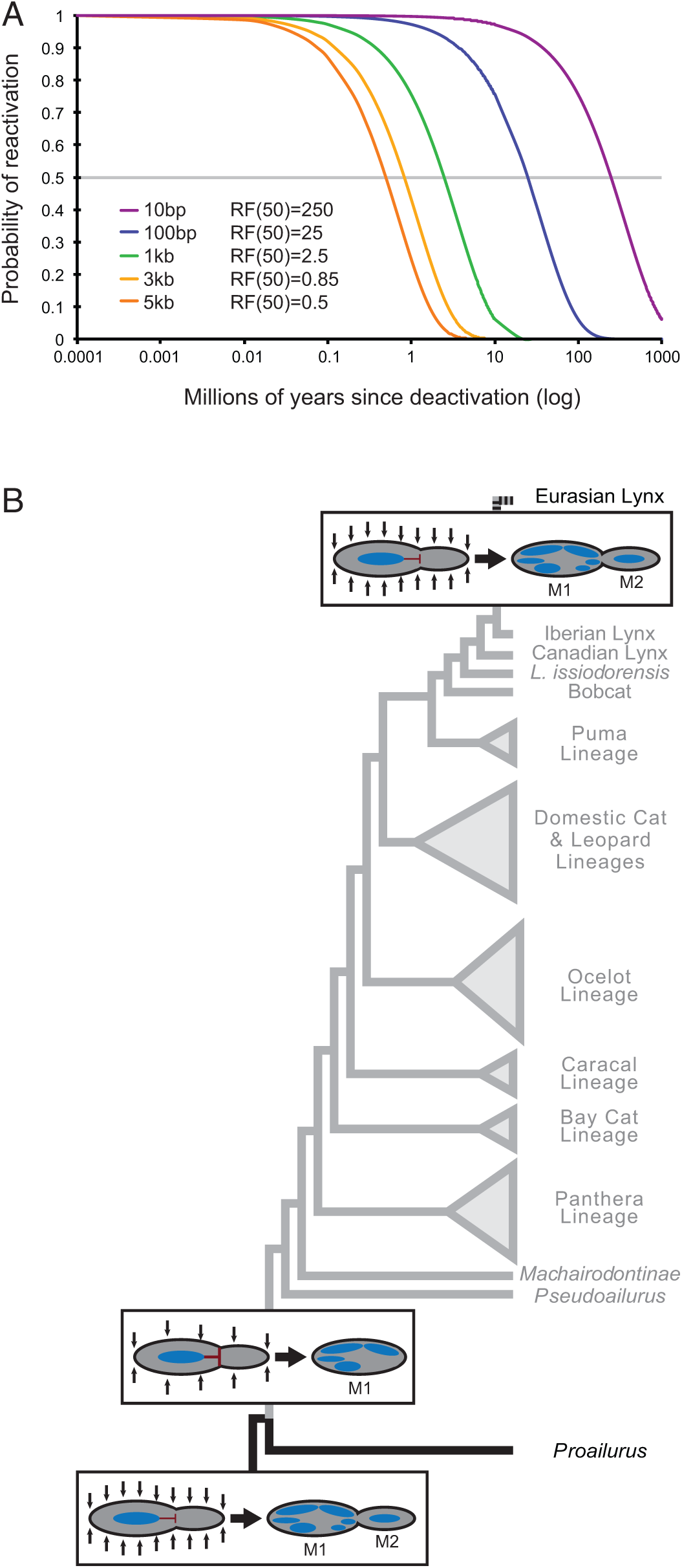
A simple model for the reevolution of lost complex traits. **(A)** The probability that functional DNA units, such as genes, regulatory elements, Transcription regions factor binding sites, and encoding protein-protein interactions within genes, can be reactivated after they are deactivated. Note that smaller elements have a greater probability of reactivation. **(B)** A developmental model for the loss of M_2_ in cats and reevolution of M_2_ in Eurasian lynx. Molar number is regulated by an activator– inhibitor model in the molarization field (black box), in which an inhibitory signal from the first developing molar (M_1_) can inhibit (red blunt arrow) the development of M_2_ and mesenchymal activation signals (black arrows) can promote the formation of molars distal to M_1_. Molar number can be reduced from two to one in felids (gray lineages) through an increase in the ability of M_1_ to inhibit (large red blunt arrow) M_2_ and/or a decrease in mesenchymal activation (fewer black arrows) leading to the loss of M_2_. Reevolution of M_2_ in Eurasian lynx may have occurred by a reversal of this developmental pattern such that there is a decrease in the ability of M_1_ (small red blunt arrow) to inhibit M_2_ and/or an increase in mesenchymal activation (more black arrows) leading to the emergence of M_2_.

### A simple developmental model of M_2_ reevolution in Eurasian lynx

The evolution and development of mammalian dentition has been intensely studied, leading to the generation of experimental- and developmental-based models that can predict patterns of mammalian tooth development. The development of mammalian molars, in particular, has been well-studied including the timing of molar initiation, mineralization, eruption, molar size, and number (Kavanagh et al., 2007). Mammalian molars, for example, develop sequentially in an anterior to posterior direction, resembling the development of serially homologous segmental structures. Kavanagh et al (2007) have described an activator–inhibitor model of sequential tooth development that describes the relative size and number of molars in mouse. In their model, an inhibitory cascade determines molar size differences along the jaw, such that the M_2_ always makes up one-third of total molar area, and that the first developing molar (M_1_) could inhibit the development of subsequent molars including M_2_. Furthermore, the initiation of posterior molars depends on previous molars through a balance between intermolar inhibition and mesenchymal activation through BMP4 and Activin A (which are highly expressed in the mesenchyme at the onset of molar formation).

These data suggest that molar number was reduced from two to one in felids through an increase in the ability of M_1_ to inhibit M_2_ and/or a decrease in mesenchymal activation leading to the loss of M_2_ (**Figure 4B**). Thus, the reevolution of M_2_ in Eurasian lynx may be a relatively simple reversal of this developmental pattern such that there is a decrease in the ability of M_1_ to inhibit M_2_ and/or an increase in mesenchymal activation leading to the emergence of M_2_ (**Figure 4B**). Remarkably Kurtén (1953) suggested that two processes, limits to the modification of existing features (i.e., evolutionary constraints) and linked development of phenotypic traits, may explain the reevolution of M_2_ in Eurasian lynx. Specifically, Kurtén (1953) suggested that selection for the enlargement of M_1_ in Eurasian lynx, which is significantly larger in than other species, may be limited but reach a size that, through developmental linkage, raise M_2_ above the threshold of phenotypic expression. Werdelin (1987) elaborated on this model to suggest that selection pressure for an enlarged molar region was greater in Eurasian lynx than enlargement of M_1_ alone could accommodate, and M_2_ reached the level of phenotypic expression through developmental linkage with M_1_ in the molarization field (Butler 1939). Thus Kurtén and Werdelin proposed a extraordinarily prescient model for the reevolution of M_2_ in

Eurasian lynx long before the establishment of mathematical and developmental models of mammalian dentition.

## Conclusions

The long half-life of enhancers, transcription factor binding sites, and protein-protein interaction motifs suggests that evolutionary reversals are possible after much longer periods of loss than previously suspected. For example, the Cypriniform oral enhancer of *Dlx2b* has retained its ability to drive proper *Dlx2b* expression in oral teeth more than 50 millions after the loss of oral teeth in Cypriniforms indicating it has escaped from loss of function mutations (Jackman and Stock 2006). More remarkably, the rudiments of the developmental program leading to tooth formation are maintained in birds 70-80 million years after the loss of avian teeth (Harris et al., 2006). Thus, components of developmental programs can be maintained for extremely long periods of time in the absence of selection acting on an expressed structure, providing a starting point for the reemergence of lost characters. Similarly, shared developmental programs can maintain the developmental genetic information necessary for previously lost characters, such as M_2_ in Eurasian lynx, to reevolve even after long periods of absence (in this case ∼15 million years).

## Acknowledgements

VJL would like to thank G.P. Wagner (Yale University) for discussions on an earlier version of this manuscript.

## Author Contributions

VJL conceived and designed the study, analyzed the data, drafted the manuscript and figures.

